# Genome-scale reconstructions to assess metabolic phylogeny and organism clustering

**DOI:** 10.1101/2020.10.07.329516

**Authors:** Christian Schulz, Eivind Almaas

**Affiliations:** Department of Biotechnology and Food Science, NTNU - Norwegian University of Science and Technology, Trondheim, Norway; K.G. Jebsen Center for Genetic Epidemiology Department of Public Health and General Practice, NTNU - Norwegian University of Science and Technology, Trondheim, Norway

## Abstract

Approaches for systematizing information of relatedness between organisms is important in biology. Phylogenetic analyses based on sets of highly conserved genes are currently the basis for the Tree of Life. Genome-scale metabolic reconstructions contain high-quality information regarding the metabolic capability of an organism and are typically restricted to metabolically active enzyme-encoding genes. While there are many tools available to generate draft reconstructions, expert-level knowledge is still required to generate and manually curate high-quality genome-scale metabolic models and to fill gaps in their reaction networks. Here, we use the tool AutoKEGGRec to construct 975 genome-scale metabolic draft reconstructions encoded in the KEGG database without further curation. The organisms are selected across all three domains, and their metabolic networks serve as basis for generating phylogenetic trees.

We find that using all reactions encoded, these metabolism-based comparisons give rise to a phylogenetic tree with close similarity to the Tree of Life. While this tree is quite robust to reasonable levels of noise in the metabolic reaction content of an organism, we find a significant heterogeneity in how much noise an organism may tolerate before it is incorrectly placed in the tree. Furthermore, by using the protein sequences for particular metabolic functions and pathway sets, such as central carbon-, nitrogen-, and sulfur-metabolism, as basis for the organism comparisons, we generate highly specific phylogenetic trees. We believe the generation of phylogenetic trees based on metabolic reaction content, in particular when focused on specific functions and pathways, could aid the identification of functionally important metabolic enzymes and be of value for genome-scale metabolic modellers and enzyme-engineers.

## Introduction

Phylogenetic trees have been important in systematizing information in biology for several centuries [1]. While the mathematical construction of these trees has not changed significantly over the years, the biological basis used for generating the phylogenetic trees has changed radically, especially during the last decades and years due to the fast development in sequencing technology [1–3]. This research field is still not settled, as exemplified by the recent change from using 16/18 S rDNA to a selection of conserved (ribosomal) proteins/genes still a topic for research [2–4].

Additionally, whole genome and genome-scale data approaches allowed by the rapid development in computational methods and computation hardware are broadening the species tree among all taxa [5–7], for example by gaining an increased resolution by reducing statistical errors due to too few comparisons [8]. One rising challenge is that this may lead to an increase in systematic errors [8]: Further errors can occur since genes may mutate due to evolutionary pressures, genes with unrelated functions may introduce artifacts into the process of generating the tree of life [3], or genes can be rearranged, nucleotides substituted or even parts of genes (for example introns) lost [9]. Depending on the research question under investigation, exactly these changes and differences might be highly interesting. Currently, there exists several approaches to integrate more sequencing data into phylogeny determinations, as for example the use of whole-genome-scale phylogeny [10]. Here however, we will focus on using genome-scale metabolic reconstructions as the foundation for determining phylogenetic trees based on metabolic capability.

A genome-scale metabolic reconstruction consists of as many as possible metabolic reactions that are encoded in an organism’s genome, thus representing the organism’s capabilities by its metabolic repertoire: Based on the presence of a gene in the genome, the organism can (in theory) perform a certain biochemical reaction and convert involved metabolites. The standard genome-scale metabolic model does not include information whether the respective proteins are expressed in the organism or not. While more advanced approaches that expand these models with a variety of ‘omics data to include such information is also possible, they have only been implemented for a limited set of organisms so far [11–14].

Lineage information is highly relevant when creating genome-scale metabolic models: Particularly in the process of model curation, it provides an important aid in the identification of possible gaps using genomic information from closely related organisms [15–17]. However, lineage information is also quite useful in determining potential horizontal gene transfers in models for Bacteria and fungi [10, 18, 19]; for example the sequences of *Staphylococcus aureus, Streptococcus pyogenes, Escherichia coli* K-12, and *Bacillus subtilis* indicate horizontal/ lateral gene transfers [20–23]. In this way, phylogeny can be of high relevance not only for curating a genome-scale metabolic model, but in particular for metabolic- and cell engineering.

When an organism is removed from its natural environment, adaptations will change the genes encoding active metabolic enzymes. In yeast, for example, entirely different pathways are expressed in anaerob versus aerob conditions. Traditional phylogenetic analyses would likely identify the adapted yeast strain as an unevolved yeast even after thousands of generations, since they are based on the use of highly conserved and stable genes. Due to changes in environmental selective pressures, mutations could accumulate in inactive metabolic genes [24, 25]. These mutations could render such enzymes less effective, causing a fitness disadvantage or even lethality when the organism is transferred back into the original environment [24–26]. We are then left with the curious situation where the adapted organism is considered by conserved genes to be identical to its origin, yet unable to live in its perceived natural habitat.

Mutations that allow adaptation to the new environment also occur in active genes (such as regulatory genes or genes coding for metabolic enzymes) to support evolution towards maximal growth fitness only after a few hundred generations [27–29]. Additionally, the organism could increase its metabolic repertoire through horizontal gene transfer from other organisms already present in the new environment. The additional and mutated genes may lead to artifacts in whole genome comparisons [3], potentially further complicating such analyses. In contrast to using whole-genome sequence-data or conserved genes to determine the similarity between a pair of organisms, the occurrence of a metabolic reaction depends on genome annotation, which is based on the detected level of sequence similarity. Consequently, binary comparison of presence of metabolic capability is independent of the immediate genetic sequence, and therefore, not subject to these mentioned artifacts.

The KEGG database (Kyoto Encyclopedia of Genes and Genomes [30–32]) has in the past been used as a starting point for calculating organism-distance measures that are based on metabolic capability [33]. The resulting phylogenetic tree showed good correlation with evolutionary distances between the organisms [33]. This allows for comparing distant organisms across domains and offers to improve model building and curation for distant organisms or a poor data basis [34].

When engineering a pathway for increased production of specific compounds, typically closely related organisms are consulted for genome-scale reconstruction curation, but also for selecting a source organism for inclusion of such reactions. Knowledge about a previous horizontal gene transfer can allow for easier metabolic engineering if transformations from this strain are already known to work *in vivo*. It may also provide actionable knowledge for model curation, specifically in identifying and solving gaps in pathways when not based on conserved genes but instead metabolic functionality. An example application of such a systematic approach is the computational tool *CoReCo*, which is designed to generate gap-less genome-scale metabolic models by integrating data using a probabilistic framework [34].

While methods based on tallying the presence or absence of metabolic functions can be computationally quite efficient, there is also much insight to be gained in explicitly analyzing metabolic enzyme similarities. To determine the changes and similarities among taxa, we present a method that uses purely evidence-based genome-scale metabolic reconstructions generated by *AutoKEGGRec* that only rely on the well-established KEGG database without any further curation [14]. In addition to a network-based binary comparison, we also conducted comparisons based on enzyme sequences of reactions present in each genome-scale metabolic reconstruction to investigate clustering among the organisms. This allows for highly specific comparisons based on single or a set of pathways, such as central carbon-, nitrogen-, or sulfur-metabolism, to a more non-specific whole-organism comparison using all the metabolic reactions to identify not only common reactions but conserved proteins across organisms.

## Results and Discussion

Starting from a set of 975 organisms (see Supplementary table 7 for details on the organisms) selected from the KEGG database currently consisting of 6, 758 organisms, we investigate the penetration of metabolic reactions, i.e. how is the metabolic reaction-set of an organism comprised of reactions that are unique to that organism or reactions that are used by many organisms. Note that, our selection consists of 134 Eukaryota (24.8% of all KEGG Eukaryota), 773 Bacteria (13.1%) and 68 Archaea (20.3%). Our selected set of organisms implement a total of *R*_*T*_ = 6, 154 different metabolic reactions (see Materials and Methods section for further details). If the metabolic reaction sets are dominated by common reactions, a binary presence or absence test as basis for generating organism-distance measures will provide low resolution in differentiating between organisms.

Defining a reaction present in less than 10% of the organisms as low penetration (LP), more than 90% as high penetration (HP), and presence in 35 – 65% as medium penetration (MP), we find that an average organism is composed of 7 *±* 7% LP, 25 *±* 5% MP, and 14 *±* 6% HP reactions. Note, that more than half (58%) of the reactions present in the data set of 975 organisms are counted as LP reactions (3, 571). We hypothesize that the LP reactions are associated with highly specialized biochemical functions not associated with central pathways. Similarly, it seems reasonable to assume that HP reactions are part of central pathways.

To gain a better understanding of the biochemical functions associated with the HP reaction set, we analyzed their KEGG pathway association (see Supplementary table 6 for details). Focusing on an HP subset of reactions, those present in at least 925 of the organisms, we find that these 78 reactions are associated with 176 KEGG pathways, of which 64 reactions are in the four enzymatic classes transferase (29), ligase (23), isomerase (7), and oxidoreductase (5). In Fig. 1 we give a more detailed overview of the pathway association for these 78 reactions.

**Fig 1.**
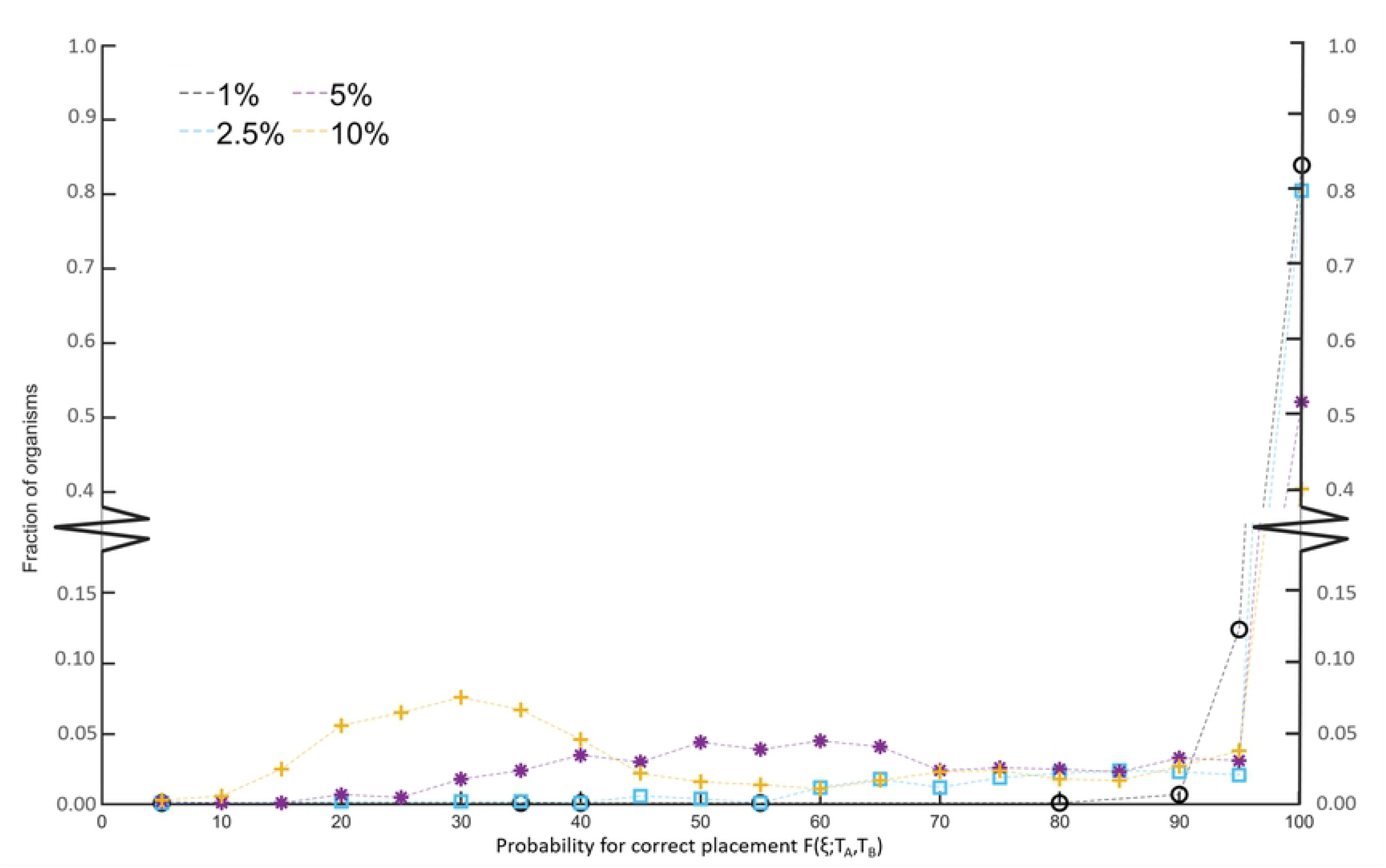
Pathway association among most frequently shared reactions. The 78 reactions found in at least 925 of the 975 organisms are associated with 176 different KEGG pathways. Note that, six of these high-penetration reactions have no pathway assignment in the KEGG database.

Note that, many reactions are associated with multiple pathways in KEGG, and surprisingly, six of the HP reactions are not assigned to any KEGG pathway. Most of the HP reactions are connected with central carbon metabolism and nucleotide- and amino acid synthesis, the central elements for living organisms, in support of our centrality hypothesis. However, we do find exceptions to this: The KEGG reactions *R03596* (937 organisms, oxidoreductase, selenocompound metabolism pathway), *R09372, oxidoreductase* (937 organisms, no assigned enzyme group, selenocompound metabolism pathway), *R04771* (945 organisms, transferase, no assigned pathway), and *R04773* (965 organisms, ligase, selenocompound metabolism and metabolic pathways) are not in central carbon metabolism pathways. These reactions all contain selenium-based-compounds. Selenium is an element appearing in Eukaryota, Bacteria, and Archaea [35–37]. Specifically, Nancharaiah and Lens [37] show a 16S rRNA phylogentic tree of selenium oxyanion-respiring microorganisms, including *Pseudomonas, Bacillus*, and *Clostridium* strains showing the diversity to their selected organisms. For the current selction of 975 KEGG organisms, selenium is an essential component for a subset of Eukaryotes and Bacteria as they are selenium-reducing. These organisms use selenium oxyanions as electron acceptors reducing it to insoluble and nontoxic selenium used in further pathways [37]. We observe however, that for most of the 975 organisms, selenium is neither essential nor significantly beneficial, yet in fact 96.1% of the organisms contain multiple selenium metabolizing reactions.

Next, we investigate the level of shared pathway association between LP and HP, finding that 2,794 (78.4%) of the LP reactions are not associated with metabolites found in HP reactions. Furthermore, 1,295 (36.3%) of the LP reactions are not assigned to any KEGG pathway at all (see Materials and Methods for details), in contrast to only 27 (18.5%) of the reactions from the HP reaction set, in support of our centrality hypothesis. These results highlight the importance of a functional-capability focus when implementing organism clustering. We therefore conclude that organism-fingerprint based phylogeny taking advantage of high-specificity (LP reactions) in combination with HP and MP reactions generates a divers yet accurate spectrum across all species.

In Fig. 2, we show the histogram of shared reactions among the 975 organisms (black symbols). We note that there is only 12 reactions that are common to all the 975 organisms, and total of 146 present in 90% or more of the organisms. In fact, the largest number of reactions are used in only a few organisms, with 375 reactions appearing in only a single organism.

**Fig 2.**
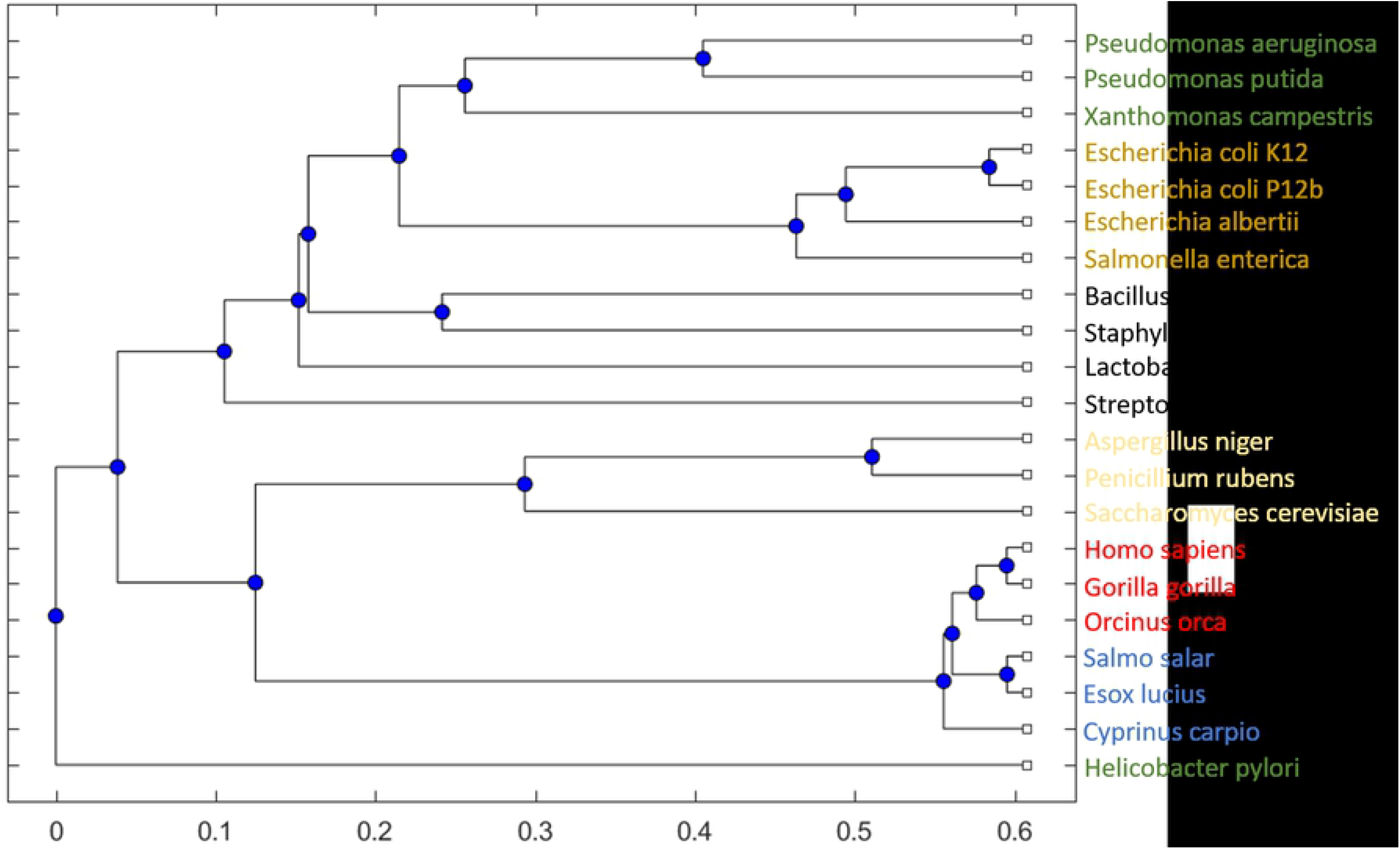
Reaction penetration in the organism set. Data from empirical networks from KEGG (black) are plotted against 10^6^ randomly generated organism reaction sets (red), together with the standard deviation curve (blue). **A)** Histogram of shared reactions (the number of organisms in which a reaction is used) among the 975 KEGG organisms with a bin size of 5. **B)** and **C)** show a magnification of panel **A)** in the LP and HP ranges, with bin size of unity.

As a contrast to the empirical data, we generate 10^6^ randomly selected reaction sets for each of the 975 organisms and calculate the expected reaction penetration in each of the random organism draws. Fig. 2 shows that the number of shared reactions for the random sets (red) is quite different from the empirical results in the low and high range:

In the empirical set, we find a significantly larger number of LP and HP reactions (panels B and C). For the random sets, we instead find that the largest number of reactions is present in approximately half of the available organisms (MP). Here, it is also quite rare for reactions to be present in only a few or many of the organisms. In Fig. 2, panel C clearly indicates the importance of shared reactions in the empirical bio-chemical reaction networks, which we will further exploit for the phylogenetic analysis.

### Fingerprint-based network comparison

We generated a phylogenetic tree using a binary metabolic network comparison approach by using a presence / absence test on all reactions for each pair of organism in the set of 975 organisms (Fig. 3, see Materials and Methods for details). As expected, we notice that the domains are separated into Eukaryota (purple), Bacteria (blue), and Archaea (green). However, we also discovered some misplaced organisms (black band) that were placed at the root of the tree, leading us to question the approaches and the stability of the network. To study the robustness of the resulting phylogenetic tree to perturbations in the particular reaction content, we conducted a sensitivity analysis by introducing a fixed level of random reactions *ζ* into each organism (see Materials and Methods for details). The results for the level of whole-tree comparisons in Tab. 1 show that the one-percent-level perturbation of the reaction network generates highly similar trees when conducting the perturbation experiment 10^6^ times (see Materials and Methods for details). Note that we use the *ζ* = 0 tree as our reference (correct placement) in the comparisons.

**Table 1.** Sensitivity analysis of phylogenetic tree. Introducing a fixed level of random reactions *ζ* results in trees that showed a similarity *K*(*ζ*) to the unperturbed tree (*K*(0)).

| $\zeta$ | 0.01 | 0.025 | 0.05 | 0.10 |
| --- | --- | --- | --- | --- |
| $K(\zeta)$ | 0.980 | 0.948 | 0.805 | 0.670 |

**Fig 3.**
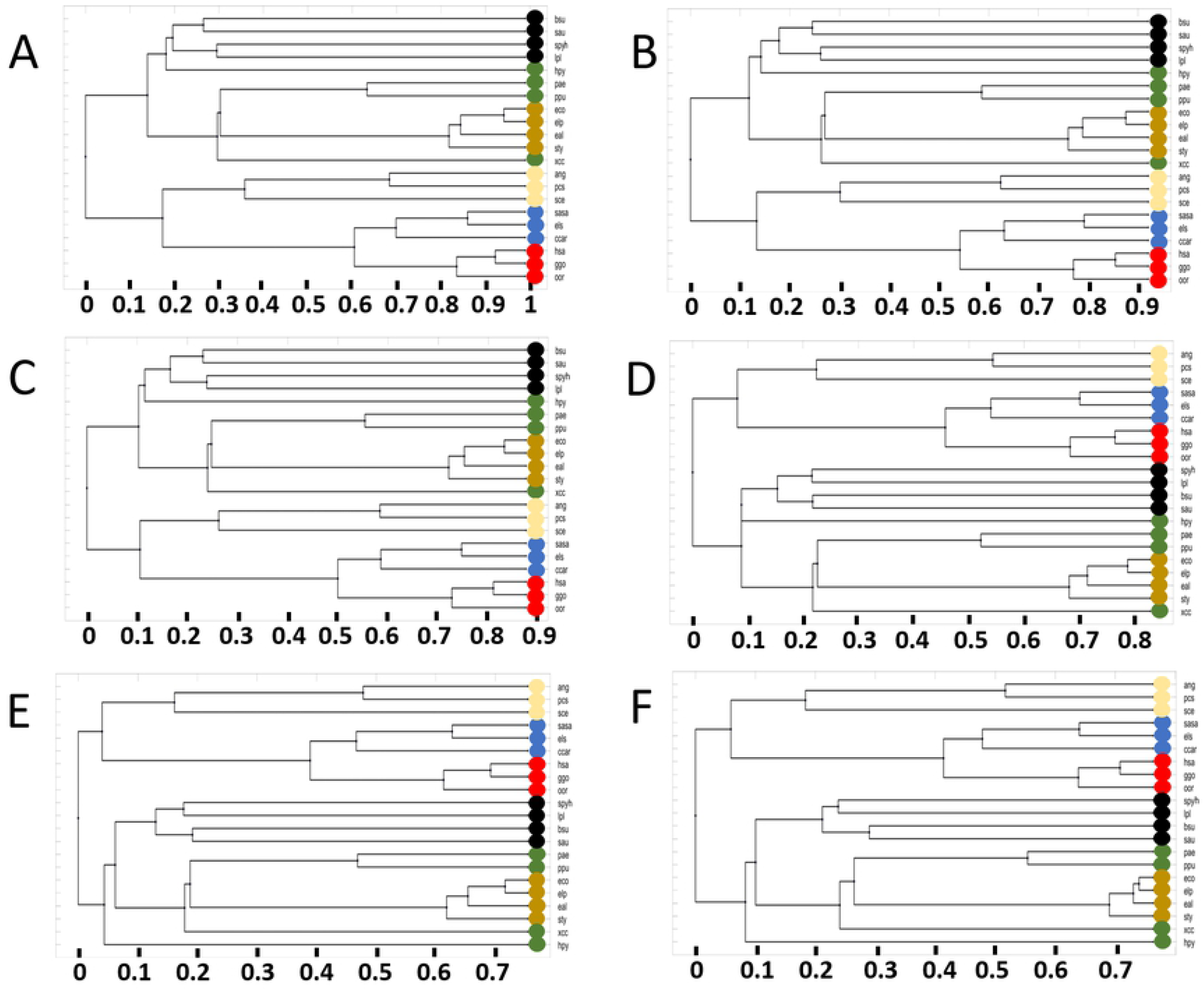
Phylogenetic tree using binary fingerprint comparison from KEGG. Using KEGG 975 organisms, the resulting tree displays a clear separation between the domains, splitting into groups such as plants, fish, birds, mammals and fungi. The red circle marks the edge of the terrabacteria group to others, such as proteobacteria. Figure created with [38].

Fig. 4 shows a histogram of the probability of correct placement of tree-neighborhoods for the 975 individual organisms. For the *ζ* = 0.01 level of randomness, we find that 83.9% of the organisms have an unchanged neighbourhood over the 10^6^ instances, and only 5 organisms show less than a 80% chance of correct placement. Increasing the level of randomness to *ζ* = 0.025, we find a slight reduction in the fraction of always correctly placed organism to 80.5%. However, increasing to *ζ* = 0.05% this has dropped significantly to 52.1%, and with *ζ* = 0.10 we find that only 40.1% of the organisms are always correctly placed.

**Fig 4.**
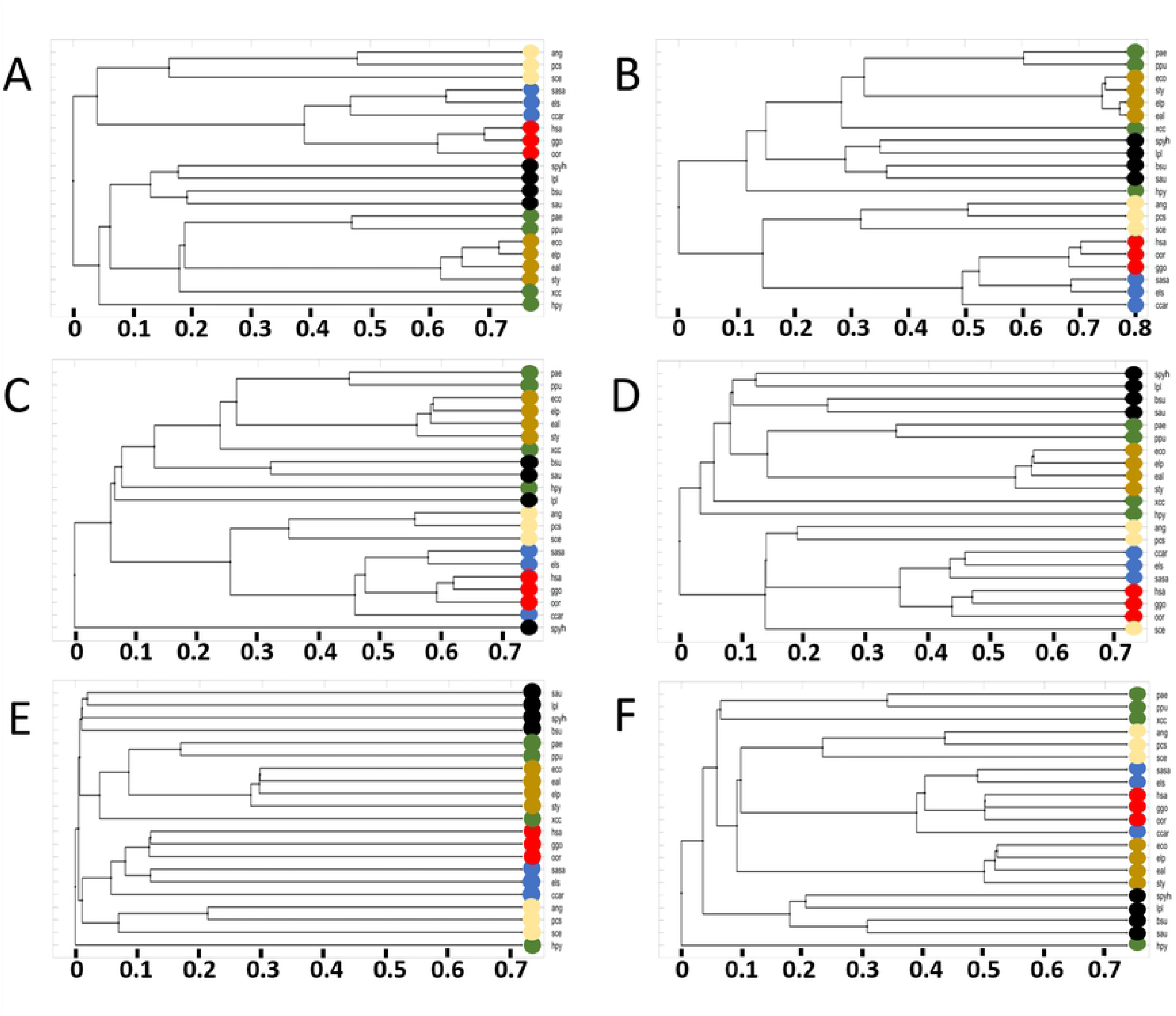
Histogram of the probability for correct placement of an organism using noisy reaction network. For all 975 organisms, 10^6^ instances or each metabolic network with 1% (black) randomized metabolic reactions, 2.5% (blue), 5% (purple), and 10% (yellow).

We continue by taking a closer look at three organisms with approximately the same number of metabolic reactions in KEGG and that are correctly placed in the *ζ* = 0 tree compared to the tree of life (see Tab. 2). Their response to increasing levels of perturbations in their reaction networks is quite dissimilar: While they respond quite similarly to *ζ* = 0.01 and *ζ* = 0.025 levels of randomness result, increasing *ζ* further has drastic effects. Already *ζ* = 0.05, which on average corresponds to a perturbation of only 76 reactions, the probability for *Xanthomonas* to be correctly placed decreases to 0.83%. For *ζ* = 0.10, this has dropped to 0.56. In stark contrast, we observe that the highly curated *Saccharomyces* reaction set is ultra stable. Even at the 10% noise level, the yeast is correctly placed 99% of the time. We argue, that this sensitivity analysis gives an indication about the quality of the reaction network.

**Table 2.** Sensitivity analysis of specific networks. Six organisms of which three are placed correctly (C) and three incorrectly (N) in Fig. 3. The organisms are specified by their KEGG codes and are compared by their sensitivity to correct placement and their reaction-penetration composition.

| Organism | Placement state | # reactions | $\zeta = 0.01$ | $\zeta = 0.025$ | $\zeta = 0.05$ | $\zeta = 0.10$ | HP | MP | LP |
| --- | --- | --- | --- | --- | --- | --- | --- | --- | --- |
| *Xanthomonas campestris* (xcc) | C | 1,520 | 0.99 | 0.97 | 0.83 | 0.56 | 9.61 | 30.9 | 5.59 |
| *Saccharomyces cerevisiae* (sce) | C | 1,523 | 1.00 | 1.00 | 0.99 | 0.99 | 9.59 | 19.9 | 11.5 |
| *Bacillus subtilis* (bsu) | C | 1,518 | 0.99 | 0.98 | 0.95 | 0.81 | 9.62 | 27.4 | 5.67 |
| *Lokiarchaeum* sp. (loki) | N | 845 | 1.00 | 1.00 | 0.76 | 0.56 | 15.6 | 26.1 | 12.1 |
| *Borrelia burgdorferi* (bbu) | N | 309 | 1.00 | 0.98 | 0.56 | 0.21 | 29.5 | 19.7 | 0.32 |
| *Candidatus Hodgkinia cicadicola* (hcc) | N | 71 | 1.00 | 1.00 | 1.00 | 0.65 | 32.4 | 21.1 | 0.00 |

Furthermore, the correct placement of a specific organism depends on the network size and the percentage of miss-annotated reactions or missing/added reactions: It is to be expected that a selection of 975 organisms across all domains from the (currently) 6, 758 organisms in the KEGG database will include some organisms with a low number of annotated reactions. Indeed, some of the incorrectly placed organisms in Fig. 3 are *“Candidatus”*-organisms that have been characterized but not yet been cultured. The set of 975 organisms contains 37 *Candidatus* strains, of which 17 are placed incorrectly: Not only are they outside of the expected groupings, but instead they are placed near the root of the tree. Consequently, they contain few reactions that are able to generate close similarity with other organisms. The *Candidatus* strains make up almost half of the 35 misplaced organisms in Fig. 3, highlighted with a black arc. However, the incorrectly placed strains are only associated with an average of 360 *±* 190 metabolic reactions in the KEGG database, in contrast to the average of 1, 200 *±* 500 for the 940 correctly placed organisms (for details see Supplementary table 8). An example of a incorrectly placed *Candidatus* strain is KEGG ID *hcc* with the name *Candidatus Hodgkinia cicadicola* TETUND1. This is an entry with only 71 reactions, of which 0% are LP, 21.1% MP, and 32.4% HP (see Tab. 2).

On the opposite end of the spectrum among incorrectly placed organisms is the organism with the KEGG ID *loki*, an Archaea within the Asgard group according to taxonomy, a *“composite genome assembled from a metagenomic sample”* from the Arctic Mid-Ocean Spreading Ridge, located 15 km from Loki’s Castle active vent site [39]. It contains 845 reactions, 12.1% of which are LP type, 26.1% MP, and 15.6% HP type, and thus, it is within the expected distribution among LP / MP / HP reactions (see Tab. 2). Note however, 7.9% of its LP reactions are only present in less than 1% of the organisms. These highly specific and unique reactions decrease the potential overlap with other species which would allow for a similarity-based placement within the tree. We find it reasonable to assume that these strains are missing a large number of metabolic reactions while also containing few strain-representative metabolic reactions, making a high quality binary reaction-presence comparison difficult. Consequently, we cannot expect to place them correctly in the phylogenetic tree.

Since most of the organisms contain a balanced mixture of LP, MP and HP reactions, the whole-metabolism comparison is capable of generating a clear separation between Eucaryotes and Procaryotes, and within the domains, a clear separation into families. In Fig. 3 the organism clustering is indicated by colored bands, and the domains Archaea (green), Eukaryota (purple), and Bacteria (blue) are highlighted separately. Within Eukaryota, the phyla are sub-grouped as expected, all fish are placed within actinopterygii (ccar, sasa) and are separated from mammalia (hsa, ggo), within the subgroup of tetrapoda, also placing sauropsida with the selected organisms correct. Next to mammalia, nearly equidistant, are viridiplantae before fungi, where aspergillus (ang) and penicillium (pcs) cluster as expected. Within Archaea, the euryarchaeota are correctly split into stenosarchae and other organisms present in the 975-organism data set.

The Bacteria domain is with 773 (79.3%) of the organisms the largest part in our data set. The gram-positive lactobacillaceae - and streptococcaceae-family (lpl and sphy) separate as expected and cluster with closely related organisms within the class of bacilli, where a correct placement of the staphylococcaceae- and bacillaceae-family (sau and bsu) within the bacillales order is achieved. The terrabacteria group is clearly distanced from the other bacteria, marked by a red circle in Fig. 3. The first class in proteobacteria that separates is alphaproteobacteria, followed by the large group of gammaproteobacteria. Separation into xanthomonadales (xcc), pseudomonadales (ppu and pae), oceanospirillales, and enterobacterales (eco and eal), to name a few, is achieved in high accordance with phylogenetic trees based on conserved genes [40]. Thus, conducting a binary finger-print comparison based on whole-genome metabolic reaction content generates a reliable tree with high accuracy and is computationally fast.

However, if we are interested in a function-based comparison between a set of organisms instead of a whole-genome comparison, a more detailed comparison including sequence information is needed: First, we can expect that focusing on a small subset of metabolic reactions will render many pairwise comparisons as either a perfect or a zero match, thus being unable to reasonably discriminate between them. Second, genes transferred into a broad-host-range plasmid can reach not only closely related organisms. This allows for fast genetic drift and numerous genotypes since, e.g. organisms sharing an environment will have a chance of sharing genetic material, and eventually, some of this will encode for metabolic reactions that are beneficial in this environment [18]. The binary network comparison will not change notably due to such processes (see Eq. (1)). Similarly for sequence comparisons that take all metabolic reactions into account. We propose that a possible solution to the challenge of conducting a narrow function-based comparison with high discriminatory power is to compare the protein sequences of selected reactions from the draft metabolic reconstructions. In the following, we report the result from such an analysis.

### Protein-based network comparisons

We start by selecting a subset of 21 selected organisms from different regions in the phylogenetic tree (see Fig. 3) to serve as our reference set. By limiting the analysis to a smaller set of organisms, it is more straightforward to study the direct consequences of different versions of the comparison algorithm and to assess consequences of a narrow function focus. Additionally, the computational run-time is significantly faster for sequence comparisons of 21 organisms versus 975. We re-generate the binary-reaction set comparison-tree for the 21-organism reference set based on Eq. (1) to serve as a gold-standard for the sequence-based analysis with focus on specific (small) sets of metabolic functions. In Fig. 5, we see that the separation between domains and even the smaller groupings can be achieved with a high degree of agreement with the 975-organism tree (Fig. 3). Most of the color coded clusters are grouped as in the large phylogenetic tree; even though small differences can be observed. Note that, when comparing the details of Figs. 3 and 5, any observed difference is solely caused by the reduction in number of organisms included in the comparison.

**Fig 5.**
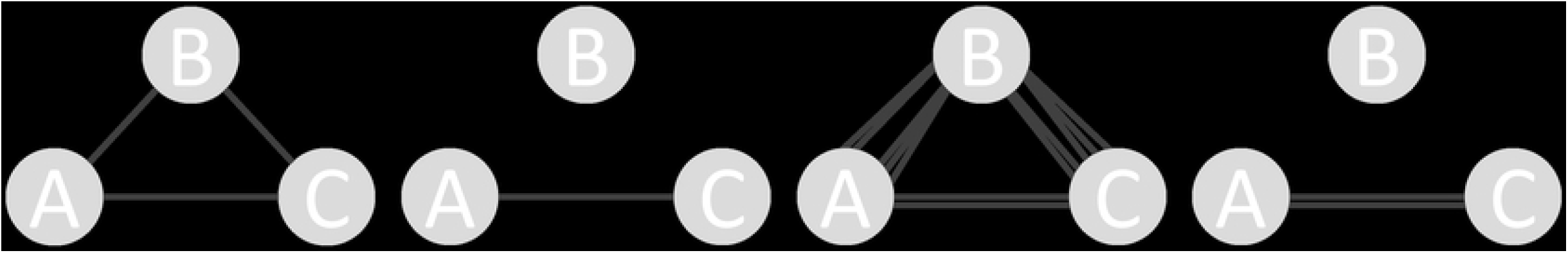
Reference phylogenetic tree using binary network-based fingerprint comparisons. For the set of 21 organisms using all reactions, the resulting tree shows clear separation between the domains. We note that the reaction network of *H. pylori* is different from the other Bacteria, and that the carp (*Cyprinus carpio*) was expected to cluster with the other fish.

As shown in Tab. 4, the number of reactions vary vastly among the set of 21 organisms: We find 2,624 biochemical reactions in *H. sapiens* but only 723 in *H. pylori*. The number of genes increases significantly, since many organisms have reactions encoded by multiple gene products through enzyme complexes or isozymes. Additionally, each of these genes may include introns or exons, silent mutations and other variations. Consequently we will compare the protein sequences associated with each enzymatic reaction that are available in the KEGG database to reduce a possible error in cross-domain-comparisons.

**Table 3.** Similarity score *K*(*A, B*) from the pairwise comparison of phylogenetic trees shown in Fig. 7.

|  | A) all reactions | B) all shared | C) TCA set | D) DNA set | E) pyruvate set | F) glycolysis set |
| --- | --- | --- | --- | --- | --- | --- |
| A) all reactions | 1.0 | 0.45 | 0.35 | 0.45 | 0.25 | 0.25 |
| B) all shared |  | 1.0 | 0.40 | 0.40 | 0.30 | 0.35 |
| C) TCA set |  |  | 1.0 | 0.35 | 0.25 | 0.35 |
| D) DNA set |  |  |  | 1.0 | 0.25 | 0.30 |
| E) pyruvate set |  |  |  |  | 1.0 | 0.20 |
| F) glycolysis set |  |  |  |  |  | 1.0 |

**Table 4.** The 21 chosen organisms in this study grouped according to order. This table also lists their KEGG IDs, number of registered metabolic reactions, number of genes, average number of genes per metabolic reaction, organism name and the associated coloring scheme.

| Order | ID | No. reactions | No. genes | average genes per reaction | organism | color |
| --- | --- | --- | --- | --- | --- | --- |
| Mammalia | hsa | 2624 | 9672 | 3.69 | *Homo sapiens* | red |
|  | ggo | 2603 | 9949 | 3.82 | *Gorilla gorilla gorilla* | red |
|  | oor | 2569 | 7984 | 3.11 | *Orcinus orca* | red |
| Neopterygii | sasa | 2504 | 17553 | 7.01 | *Salmo salar* | blue |
|  | els | 2507 | 9613 | 3.83 | *Esox lucius* | blue |
|  | ccar | 2404 | 17255 | 7.18 | *Cyprinus carpio* | blue |
| Fungi | ang | 1965 | 4942 | 2.52 | *Aspergillus niger* CBS 513.88 | yellow |
|  | sce | 1522 | 3012 | 1.98 | *Saccharomyces cerevisiae* S288c | yellow |
|  | pcs | 1931 | 4288 | 2.22 | *Penicillium rubens* Wisconsin 54-1255 | yellow |
| Proteobacteria | pae | 1716 | 3244 | 1.89 | *Pseudomonas aeruginosa* PAO1 | green |
|  | ppu | 1606 | 2866 | 1.78 | *Pseudomonas putida* KT2440 | green |
|  | xcc | 1523 | 2640 | 1.73 | *Xanthomonas campestris* pv. *campestris* ATCC 33913 | green |
|  | hpy | 723 | 925 | 1.28 | *Helicobacter pylori* 26695 | green |
|  | eco | 1743 | 2737 | 1.57 | *Escherichia coli* K-12 MG1655 | orange |
|  | elp | 1714 | 2743 | 1.6 | *Escherichia coli* P12b | orange |
|  | eal | 1610 | 2571 | 1.6 | *Escherichia albertii* KF1 | orange |
|  | sty | 1634 | 2623 | 1.61 | *Salmonella enterica* subsp. *enterica* serovar Typhi CT18 | orange |
| Bacilli | sau | 1131 | 1660 | 1.47 | *Staphylococcus aureus* subsp. *aureus* N315 | black |
|  | bsu | 1520 | 2408 | 1.58 | *Bacillus subtilis* subsp. *subtilis* 168 | black |
|  | lpl | 993 | 1684 | 1.7 | *Lactobacillus plantarum* WCFS1 | black |
|  | spyh | 760 | 1049 | 1.38 | *Streptococcus pyogenes* HSC5 (serotype M14) | black |

From the KEGG pathways, we first used all available reactions in our 21-organism reference set (see Tab. 4 for details). Due to the possibility that a reaction only appears in one of the organisms in the pairwise comparison, we found it necessary to add a penalty score (see Materials and Methods section for details). We found it further necessary to implement a solution for multiple enzymes in a single reaction, either isozymes or enzyme clusters, as a basis for the pairwise biochemical reaction-comparison. In the following two sections, we discuss the reasoning for and impact of the respective methodologies.

### Impact of penalty score on the phylogenetic trees

Pairwise binary reaction-comparison result in either *true* or *false* responses. However, when implementing a sequence-based comparison, we are interested in the possibility of a more graded response to cases where a reaction is only present in one organism. A penalty score is in essence the application of a low similarity score for such cases. This penalty score affects very similar organisms that (a) share many reactions, and (b) have a high sequence similarity the least, similar to organisms that share (a) few reactions with (b) a low similarity score.

We conduct our assessment of possible penalty scores using all available reaction-based enzyme sequences for each pair of organisms (Fig. 6). We find that a clear separation between Eukaryota and Bacteria could be achieved for all values of the penalty score. To evaluate the impact on highly similar organisms, we focus on the two *E. coli* strains and the *E. albertii* strain, with KEGG organism IDs *eco, elp*, and *eal* respectively. They are all encoded in orange colors in the 21-organism tree figures. The relative distance between *eco, elp* and *eal* do not change when varying the penalty score: As expected, *eco* and *elp* are always closer related in comparison to *eal*. The value of their similarity score, ⟨*S*(*A, B*) ⟩ (see Eq.(7)) changes, which for closely related organisms means a small change in distance in the tree.

**Fig 6.**
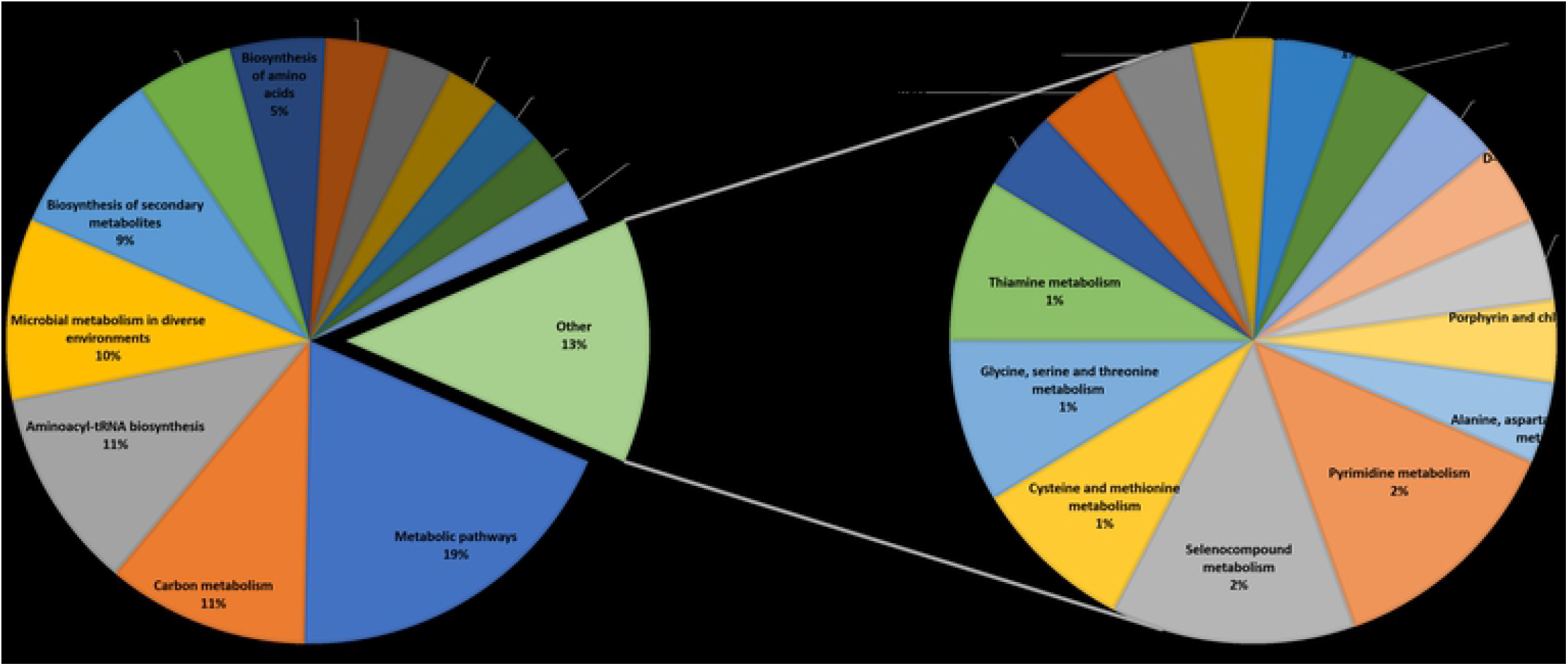
Effect of penalty score using all metabolic reactions in the 21 genome-scale networks. The different panels refer to the penalty score used: (A) − 0.25, (B) − 0.1, (C) 0, (D) 0.1, (E) 0.25 and (F) no use of penalty. The color codes and KEGG IDs of the organisms are shown in Tab. 4.

However, for distantly related organisms the penalty score has a larger impact due to the nature of Eq. (7). This is clearly evident for the green-labeled part of the phylum of proteobacteria. The organisms in the phylum proteobacteria are labeled green and orange, orange for the included family of enterobacteriaceae. While the *Pseudomonas* strains ppu and pae (green) cluster together with small differences in the tree distance, the response to varying penalty scores for the more distant *Xanthomonas* (xcc) and *Helicobacter* (hpy) does not follow a clear pattern. For a penalty score of 0.25, we find hpy clusters outside of Bacteria, whereas for a range of values of − 0.25 to 0 it’s grouped with bacilli (black). At a penalty score of 0.1, hpy is similarly distant to all Bacteria. When ignoring the use of a penalty (see Fig. 6 F), hpy clusters again outside of Bacteria. This shows that, for very close organisms the impact of the penalty score is potentially smaller, especially when few reactions are compared.

Therefore, depending on the question asked, we believe it prudent to (a) either ignore the use of a penalty, or (b) use a penalty score that has minimal impact on the similarity score between closely related organisms while at the same time modifies cluster assignments among more distantly related organisms to be more aligned to “*true functional similarity*”.

Taking these considerations into account, we found a penalty score of 0.25 be a good choice for the 21-organism sequence alignment analysis. The reasoning is based on the fact that, we achieve a high similarity with the tree based on conserved genes, even though hpy is placed away from the other proteobacteria and barely more similar to bacilli. However, the overall distances are more reasonable when comparing distantly related organisms. This is exemplified with the black labeled bacilli being more similar at a penalty score of 0.25. Additionally when focusing on specific functionality, we are not generating trees with the intent of comparing them to those resulting from conserved genes-comparisons. Instead, our reference tree is that in Fig. 5.

### Multiple enzyme comparisons for a single reaction

Even though similarities at the whole-organism level (whole genome comparisons) are of high interest, specific functional similarity is of high relevance in e.g. the field of metabolic engineering. Sequence similarities for a narrowly selected set of metabolic reactions, functionalities, and pathways can generate important insight into complex relationships based on horizontal gene transfer or mutations. Many reactions included in a genome-scale metabolic network are encoded by multiple genes where the enzymes are isozymes or complexes catalyzing the metabolic reactions.

For the cases with several genes encoding a single metabolic reaction, we generated all pair-wise sequence comparisons between two organisms (see Materials and Methods section for details). The largest similarity score, independent of sequence size or number of sequences present, was selected. While this approach could lead to a high similarity score caused by a small part of a protein complex (possibly not even part of the active site), the final similarity score between two organisms is based on the average of all protein similarity comparisons (Eq. (7)). Thus when comparing many reactions, the impact of the alignment-score from a single protein is smaller than in case of just a few reactions. Since only the highest similarity is used, we argue that even small amounts of reactions generate accurate measurements.

### Whole-genome metabolic protein sequence similarity tree

The phylogenetic tree resulting from a similarity calculation using our 21-organism reference set based on sequence comparisons is shown in Fig. 7A. It includes all reactions present in the genome-scale metabolic network with a penalty score of 0.25. To further facilitate comparisons with the corresponding binary network-based fingerprint (Fig. 5), we have provided a side-by-side rendering in Supplementary figure 1. In Fig. 7A, Eukaryota and Bacteria separate as expected when using the metabolic enzyme sequence comparison for the whole metabolic network. In contrast to the binary network fingerprint (Fig. 5), the selected *E. coli* strains share a higher similarity than salmon and pike, including *E. albertii*. The reason is that essential genes in bacteria are more conserved than nonessential genes [18, 41]. This illustrates the power of the approach utilizing potentially shared protein sequences for metabolic similarity. The higher reproduction cycle of microorganisms, including the possibility of exchange of genetic material and thus functionality from other organisms in the same environment, allows for a fast adjustment to new or changing environments. The conserved essentiality is also expressed by the fact that many metabolic reactions are encoded by a single gene in Bacteria, in contrast to more complex organisms which encode metabolic functionality often by multiple genes, compare Tab. 4.

**Fig 7.**
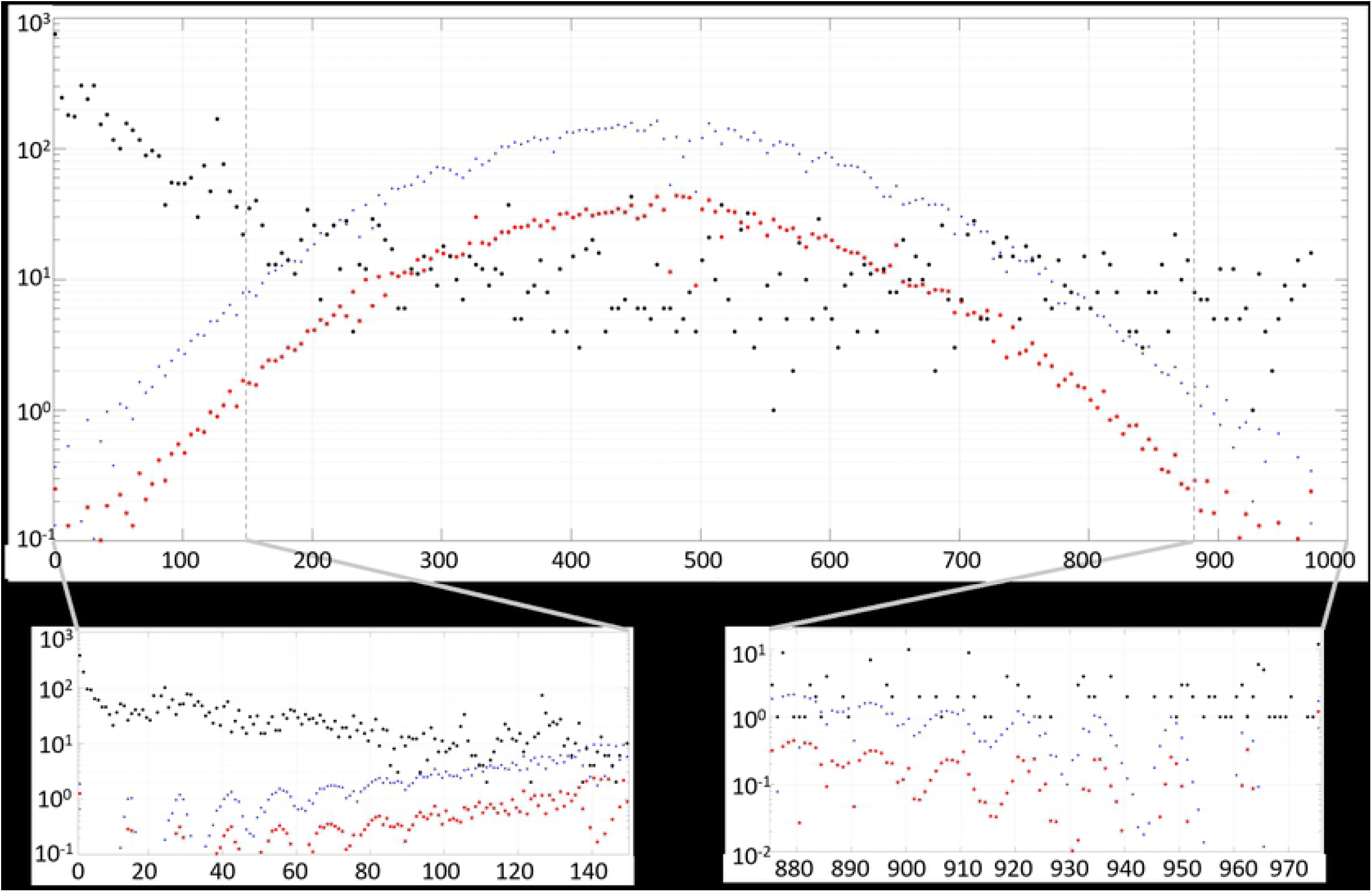
21 organisms compared by different reactions present in the genome-scale metabolic reconstructions. **A)** is a comparison based on all reactions in the network, **B)** refers to all shared reactions, **C)** is the TCA set, **D)** is based on the DNA metabolism set, **E)** compares the pyruvate metabolism set, and **F)** is with the glycolysis set. The color code and the KEGG IDs of the organisms are shown in Tab. 4, the KEGG reaction IDs are listed in the Supplementary material.

All strains meet the expected distance, even within the proteobacteria, where the orange marked family of enterobacteriaceae clusters in close proximity. Proteobacteria and bacilli are as distant as expected, however *H. pylori* clusters outside yet still within Bacteria. Note, that hpy has the smallest reaction network in this dataset of 21 organisms, reducing possible reaction overlap with other organisms. *Streptococcus* is with its 760 reactions similar in size, yet the metabolic overlap with the other organisms is larger (see Fig. 6 for more details).

Consequently, we argue, that including a higher resolution using active enzyme sequences improves the computationally fast and good functioning approach of binary metabolic network fingerprinting further. Not surprisingly, however, comparing all sequences for all organisms is significant more computational demanding.

### Consequence of specific metabolic functions on the phylogenetic trees

While comparing the whole genome-scale metabolic network on a protein sequence-basis can reveal similarities on an organism-level, we are particularly interested in similarities of metabolic reaction subsets i.e. implementation of metabolic function. This includes organism-specific functionality or analysis of conserved pathways such as the TCA cycle. As a proof of concept, here we have selected several sets of metabolic reactions based on the HP data set (see Materials and Methods section for details on the selected reaction sets). Note, that their selection is inspired by the HP data set, but driven by the KEGG pathways making some of the reactions absent in all organisms. We use the penalty score of 0.25 which imposes a high dissimilarity (penalty) on the respective missing reaction.

In Fig. 7, panels B to F, we show the phylogenetic trees resulting from the selected reaction sets. In panel B we used the protein sequences for the set of 203 metabolic reactions shared between **all** of the 21 organisms, which represents 28% of the *H. pylori* metabolic reactions, further emphasizing the importance of these reactions. The figure shows that the orca whale is closer related to the human than to the gorilla.

Furthermore *E. coli* K12 MG1655 and *S. enterica* are more similar than *E. albertii* and *E. coli* P12b, in sharp contrast to previous findings and the common understanding of the Tree of Life. Although the similarity among mammals is lower than expected using traditional comparisons, this similarity analysis is solely based on the protein sequences of the 203 KEGG metabolic reactions shared in the 21-organism reference set. Here, the choice of penalty score has no effect, since all the reactions and therefore at least one protein sequence, is available for all pair-wise sequence alignments.

Using protein sequences from metabolic reactions present in the TCA cycle (Fig. 7C), we find that the separation between Bacteria and Eukaryota is disturbed by *S. pyogenes* (spy), which is the second smalles organism with its 760 metabolic reactions. Within the Bacteria, only the family of enterobacteriaceae clusters as expected. The TCA set also generates high relative similarity between human and gorilla. The relative distance to the orca whale thus is more in accordance with traditional clustering.

In Fig. 7D, we show the phylogenetic tree resulting from the protein sequence comparison of a set of DNA-metabolism reactions, also based on the HP reaction set. This reaction set also contains all reactions shared among all 975 organisms. While we observe that all distances increase, the relative sub-groupings do not change as Bacteria and Eukaryota are separated as expected. The clade of euteleostomi as well as the family of enterobacteriaceae and class of bacilli form clusters, which are as expected with the exception that salmon and carp switched their relative similarity to pike. Note that, the other organisms in the class of gammaproteobacteria (xcc and hpy) are not placed with the other gammaproteobacteria but instead share similarities with all Bacteria.

In contrast when we use a metabolic reaction set based on pyruvate metabolism (Fig. 7E), we find the largest relative dissimilarity of all the compared reaction sets (panels B-F). Surprisingly, the expected determination of domains is still achieved. The phylogenetic tree displayed in panel F is based on a set of only seven metabolic reactions from glycolysis that are present in most metabolic networks. Similar to panel E, the relative distance of all organisms is high compared with the other reaction sets. Note that, in several of the figure panels the distance of multiple organisms are nearly identical, e.g. in panel 3 for *Escherichia* and the mammals. This result suggests that that the protein sequences for these organisms in the chosen metabolic reaction sets are equally distant to each other.

For a more quantitative analysis of the phylogenetic tree similarities, we calculated the similarity *K*(*A, B*) between all pairs of phylogenetic trees in Fig. 7 based on a branch comparing approach (further details in Materials and Methods section and Ref. [42]). We find some interesting patterns from this network comparison (see Tab. 3). Panels E and F have a low similarity (*≤* 0.35) with the other panels. Compared to the all reactions panel (A), the score of 0.25 for E and F respectively is the lowest, and panels D and B share a similarity of 0.45 with A. We conducted this tree similarity analysis as an indication for the over all organism similarity to evaluate large dissimilarities. These dissimilarities show reaction sets that vary in sequence similarity that the tree is re-ordered and some organisms cluster differently.

Close organisms were expected to cluster closely, which is achieved for many groups. The smallest set of reactions compared here was seven (glycolysis set), the largest 21 reactions (TCA set). Comparing the relative distances between these two plots shows a clear difference: While the small number of reactions leads to a large difference for even very close organisms such as the two *E. coli* strains, the large set of reactions shows a very high similarity. The pairwise comparison of the two *E. coli* strains, a similarity of 0.73 in A, 0.78 in B, 0.58 in C, and 0.53 in F thus shows the clear pattern: In A all reactions are used which also includes a penalty score for the binary reaction difference. Additionally dissimilar enzyme sequences via the average organism similarity influence the scores. In panel B only reactions are compared which are present in all strains. This leads to a pure sequence similarity based comparison and thus to closer organisms within *E. coli*.

The distances between organisms in the phylogenetic trees of panels C to F is a direct result of the particular selection of metabolic reaction sub-sets, or specific metabolic functions. A consequence of smaller reaction sets is that some of the chosen metabolic functions are not or only partly present in a given organism set. The difference can be seen when comparing the pike (els) to *E. coli* K-12 MG1655 (eco) for the TCA set shown in panel C. Firstly, the organisms differ in the presence of four reactions: KEGG reaction IDs R00344, R00432, R00709, and R01197. R01197 is the only reaction present in eco but not in els. For these reactions, the penalty score is used in the comparison. A closer inspection shows that the more complex organism, els, has a larger repertoire of alternative reactions for the succinate:CoA ligase, the enzyme catalyzing this reaction.

The biochemical transformation of succinly-CoA to succinate is an important reaction in the TCA cycle and encoded via the different enzymatic reactions with KEGG reaction IDs R00405, R00432, R00727, and R10343. Reaction R00405 allows for ATP production in eco and els. reactions R00432 and R00727 are responsible for GTP and ITP respectively in the pike. In fact, they are present in all selected 21 Eukaryota. This shows the higher metabolic versatility of the Eukaryota in this central succinate:CoA ligase reaction. Reaction R10343 has no involvement of any energy equivalent and is present in neither of these two organisms, however in is included in pae, ppu and xcc. Thus, for the similarity analysis of eco and eal, this reaction has no impact. The difference of eal and eco is consequently based on the four binary reaction differences that are included with a penalty score of 0.25 and on the sequence differences for the remaining 17 reactions.

We have shown, that the analysis of selected reactions generates a set-specific similarity. We find that the similarity strongly depends on the reactions and thus varies for the reaction sets. The reaction distribution within the family of enterobacteriaceae is so similar that they cluster together in all panels in Fig. 7. We notice however, a clear increase in their relative distances (height of the branches in the phylogenetic trees). For eco and eal, this is based on the direct sequence difference and variation in reaction availability. The large difference in panel E indicates huge variation amongst all organisms for this reaction set. We find that the analysis of reaction sets together with whole-genome metabolic comparison can increase the knowledge for specific organisms. We would expect phylogenetic trees resulting from specific metabolic function comparisons to differ from the expected Tree of Life in both distance and clustering. However, these trees are not intended to serve as replacements for the Tree of Life. Instead, they have a role as knowledge basis for engineering purposes when combining reaction sets of multiple organisms.

## Conclusion

Here, we have developed and analyzed an organism-clustering method based on genome-scale metabolic capability. We used the previously published Matlab function AutoKeggRec [14] to generate 975 genome-scale metabolic draft reconstructions based on the KEGG database. This reaction set represents various phlya from all domains of life. We find a strong similarity between the binary metabolic network-based fingerprint organism-clustering and the Tree of Life, in agreement with previous studies [33]. When using all available KEGG metabolic reactions for the organisms, we would expect that the resulting phylogenetic tree should have strong similarities with the accepted Tree of Life. Incorrectly placed organisms within the phylogenetic tree share a small overlap of metabolic reactions with other organisms. That is caused by two factors: Their small metabolic reaction network due to incomplete or significantly missing reaction annotations, or a large number of highly specialized metabolic reactions combined with a small number of more common metabolic reactions. We observed a similar effect when reducing the number of organisms from 975 to 21, as some relationships were changed. When analyzing the binary genome-scale networks’ similarities, we find that high-specificity reactions (LP reactions) and the more common MP reactions generate an accurate placement of the organisms in the hierarchical clustering of organisms. Furthermore, a correct placement of an organism also depends on the presence of HP reactions, since without these the organisms tend not to be grouped within their expected domains.

Within the HP reaction set we find universal reactions in central pathways that metabolize amino acid and nucleotide based compounds besides selenium processing reactions. The HP reaction set is thus associated with an ubiquitous central metabolism across all organisms. Further analysis generating a metabolic core network based on these data, curating and gap filling them, can increase the fidelity of similarity analyses founded on genome-scale metabolic reaction information on a systems level. Additionally, it can help in the process of curating similar and new organismal metabolic networks as well as revising annotations for existing ones.

In contrast, we find that the LP reactions which are shared between only a few organisms consist of 58% of the 6, 154 metabolic reactions present in the data set of 975 organisms. Since 78.4% of these reactions do not share metabolites with HP reactions and 36.3% are not associated with a pathway in the KEGG database, we conclude that these reactions are highly specialized metabolic reactions. However, since the LP reactions are quite abundant in the data set, they play an important role in combination with the MP reactions to achieve correct placement among closely related organisms. For example, the KEGG reaction ID R00025 is present in fungi, bacilli, and in parts of the proteobacteria, not in neopterygii, mammalia, or enterobacteriaceae.

When combined with other reactions clustering for Eukaryota and Prokariota, this reaction can contribute to the difference between enterobacteriaceae and pseudomonadales on a binary level. In contrast, it will not provide the ability to differentiate between *E. coli* and *S. entericia* since it’s not present for these organisms. In order to improve the resolution of the comparisons, we implemented a protein sequence-similarity measure with the possibility for selecting smaller reaction sets and pathways composed of specific reactions.

Using the protein sequence data, we find a higher level of agreement between the resulting phylogenetic tree and the tree of life for the reduced data set. The additional data within the enzymatic reaction sequences can therefore compensate for the small overlap in diverse organisms. We found differences in clustering when applying the similarity analysis to various reaction selections based on the HP reaction set. Consequently, we hypothesize that using the protein-sequence based similarity analysis focused on specific metabolic reaction subset may aid in the identification of functionally important mutations and adaption-relevant gene transfers across species, families or even phyla.

## Materials and Methods

In the following, we describe the two different approaches employed to generate the metabolic-network based phylogenetic trees. Both methods are based on the availability of draft genome-scale metabolic reconstructions from KEGG, using the AutoKEGGRec function [14]. This function runs in Matlab and interfaces with the COBRA toolbox 3.0 [43] for genome-scale metabolic modelling. Briefly described, AutoKEGGRec will access the KEGG database using identifiers for organisms and generate genome-scale draft reconstructions based on only the genetic information. This starting point will typically be subject to automatic as well as manual curation, gap-filling and improvements based on additional information in the standard model-creation steps [17]. In this work we only used data available in the KEGG database.

Calculations and figure generations were performed in Matlab 2017b: The Matlab function*seqlinkage* was used with the default setting *‘average’* for an unweighted pair group method average (UPDMA) clustering of the pairwise comparisons [44]. To create the figures the function *plot* was used with the *Type*-setting *‘square’*.

Using AutoKEGGRec, we downloaded the full reaction table for 975 selected organisms. The provided KEGG organism codes (see Tab. 4 for 21 organisms and Supplementary Table 1 for the *N* = 975 organisms) were used as input for AutoKEGGRec, and a community draft model as well as individual draft models were generated. Furthermore, we used the AutoKEGGRec option to generate the organism-reaction-gene matrix (ORG), which is used as the knowledge-base for this work (see Ref. [14] for further details). The total number of reactions covered by the set of *N* organisms is *R*_*T*_ = 6, 154 out of 10, 995 reactions present in the KEGG database (56.2%).

Note, that the 21 organisms are largely a subset of the 975 selected KEGG organisms chosen based on distribution and their prominence. The color code given in Tab. 4 for the organisms is consistent throughout the manuscript.

## Binary metabolic network comparison

The first method for generating phylogenetic trees is based on using the organism-reaction-gene (ORG) matrix generated by AutoKEGGRec by transforming it into a binary matrix where a unit entry indicates that the given reaction is present in the organism, and a zero entry when it is absent, refer to Tab. 5. Therefor this is a whole-network comparison of the organisms, resulting in a metabolic functional similarity. We use this binary matrix as the starting point for an all-to-all pair-wise comparison between the organisms by calculating the Jaccard index [45] *J* (*A, B*) between the pair of rows corresponding to organisms *A* and *B*:

**Table 5.** Example binary representation of the transposed of the ORG matrix from AutoKEGGRec for two organisms, A and B. This matrix is used as input for the binary network based similarity calculation, the SW align data is the result of the metabolic-reaction-based protein sequence alignments.

| Organism | R01 | R02 | R03 | R04 | R05 | R06 | R07 | R08 | R09 | R10 | R11 |
| --- | --- | --- | --- | --- | --- | --- | --- | --- | --- | --- | --- |
| A | 1 | 1 | 0 | 0 | 1 | 1 | 1 | 1 | 0 | 0 | 1 |
| B | 1 | 0 | 0 | 0 | 0 | 1 | 1 | 0 | 0 | 1 | 1 |
| $S_R(A, B)$ | 0.87 | X | 0 | 0 | X | 0.93 | 0.75 | X | 0 | X | 0.89 |

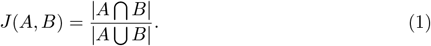

We illustrate the procedure with the following example: The similarity of the two organisms A and B is calculated following Eq. (1) and using the toy data shown in Tab. 5 as

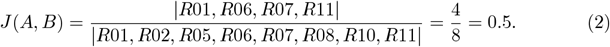

For improved calculation speed, only reactions occurring in at least one organism were included, and empty KEGG reactions were removed. The pair-wise similarity scores were stored in a matrix, saved as a Newick-formatted file, and visualized in either Matlab or iOTL [38].

### Sensitivity analysis

We investigate the sensitivity of our results to inaccuracies in reaction-associations with organisms by performing a random sampling analysis. In this way, we can assess the effect of varying levels of incorrect assignment of reactions to organisms on our conclusions, i.e. the consequence of incorrect loss and acquisition of reactions. First, we generate an instance of each network for each of the 975 organism. Second, we introduce the noise-level *ζ*. The choice of *ζ* = 0.01 means that, for each reaction in a network instance, there is a 1% chance that it is removed and a randomly (uniformly) selected reaction not assigned to the organism is chosen instead. In this way, the number of reactions for each organism stays constant. In the case of *E. coli*, the network consists of *R*_*E*_ = 1, 743 KEGG reactions. With *ζ* = 0.01, we will remove 17 of these reactions and replace them with the same number drawn from the *R*_*T*_ *– R*_*E*_ = 9, 252 remaining possible reactions. In our sensitivity analysis, we use *ζ* ∈ {0.01, 0.025, 0.05, 0.1}. Third, we conduct an all-against-all binary comparison of the metabolic networks using Eq. (1).

The topology of the phylogenetic tree *T*_*B*_ resulting from level *ζ* of randomness in metabolic reactions is compared to the reference (*ζ* = 0) tree *T*_*A*_ by counting interior branches that return the same partitions. Defining *F* (*ξ*; *T*_*A*_, *T*_*B*_) ∈ {0, 1} as a binary function that takes the value unity (zero) if the interior tree branch nearest the organism *ξ* is identical (dissimilar) in trees *T*_*A*_ and *T*_*B*_, we have

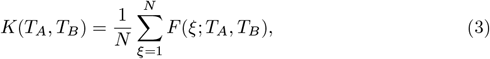

where the numeric score *K*(*T*_*A*_, *T*_*B*_) ∈ [0, 1] reflects the level of similarity between trees *T*_*A*_ and *T*_*B*_, with unity being identical. The method is described in detail in Refs. [42, 46, 47]. Briefly, the approach consists of first calculating a new rooted tree for each organism *ξ* using the *subtree* function in Matlab. The *getcanonical* function ensures a leaf sorted tree. For each leaf (organism), the local similarity is compared with the Matlab function *isequal*, resulting in the binary score *F* (*ξ*; *T*_*A*_, *T*_*B*_) for the nearest branch. This process is repeated a total of *L* = 10^6^ times for each of the chosen noise levels *ζ*, resulting in an average similarity score *K*(*ζ*) for the *L* pairwise comparisons.

When conducting the pairwise tree similarity analysis related to Fig. 7, with results reported in Tab. 3, we implemented a non-binary version of *F* (*ξ*; *T*_*A*_, *T*_*B*_): We scale each branch similarity by weight factors that are inversely proportional to the selected number of organisms. Each of the *N* − 1 branch comparisons is scaled by (*N* − *η*)*/*(*N* − 1), where *η* denotes the branch number (in increasing sequence). For a tree with four organisms this means the first branch is scaled by (4 − 1)*/*3, the second closest by (4 − 2)*/*3, and the last is scaled by (4 − 3)*/*3. Consequently, we define

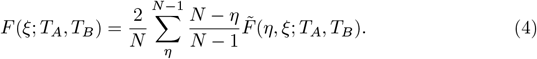

Note that for this situation, *F* (*ξ*; *T*_*A*_, *T*_*B*_) ∈ [0, 1], whereas 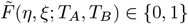 is a binary function. This ensures higher impact of local (dis)similarity in the respective comparisons.

### Shared pathway association within LP content

AutoKEGGRec provides all data linked for a molecule or reaction in KEGG for the models. This includes pathway IDs and links other data base besides the chemical reaction required for the genome-scale reconstruction. We used the chemical reaction linked by the reaction ID to identify KEGG compound IDs in the LP, MP, and HP reaction set. This allows for a fast and easy comparison for these sets, and an association of a compound to another reaction set. For the pathways linked to reactions we use the reaction IDs to scan whether they have pathway information stored.

### Alignment of metabolic-reaction protein sequences

Our second approach is based on aligning the amino-acid sequence(s) corresponding to the KEGG-coded reactions for all pairs of organisms. The enzyme(s) catalyzing each reaction of each organism according to KEGG were stored within the ORG matrix. In Tab. 5, each entry of unity is replaced by the protein sequences linked to this reaction. For each KEGG reaction *R*, the sequences were aligned using the *swalign* function in matlab, which performs a local alignment using the Smith-Waterman (SW) algorithm [48] with the default scoring matrix BLOSUM50 for proteins. The *swalign* function generates the locally aligned sequences and a score as output which was normalized using the SW score of each protein against itself. Consequently, percentual similarities of the protein sequences were generated as an alignment score:

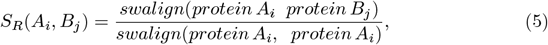

with *A*_*i*_ and *B*_*i*_ being individual protein sequences associated with reaction *R* in the two organisms, and we define the total alignment score *S*_*R*_(*A, B*) for reaction *R* as

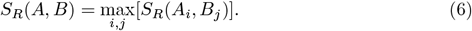

Further, we define the average comparison score as

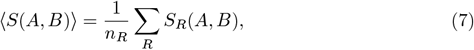

where *n*_*R*_ is the number of reactions included in the comparison between organisms *A* and *B*.

An example calculation is shown in Tab. 6, where the two organisms *A* and *B* are compared based on the respective protein sequences of their metabolic reaction R01. For this reaction, each protein of organism A has been locally aligned and normalized to each protein of organism B, resulting in an all-against-all similarity score matrix for the protein sequences associated with R01 in the two organisms.

**Table 6.** Example of protein sequence alignment of the shared metabolic reaction R01 of the two organisms A and B. In the two organisms, the metabolic function of R01 is linked to three and four proteins. The Smith-Waterman algorithm with the BLOSUM50 scoring matrix was used for the pairwise sequence alignment.

| | Protein $B_1$ | Protein $B_2$ | Protein $B_3$ | Protein $B_4$ |
| --- | --- | --- | --- | --- |
| Protein $A_1$ | 0.87 | 0.48 | 0.81 | 0.83 |
| Protein $A_2$ | 0.65 | 0.55 | 0.23 | 0.78 |
| Protein $A_3$ | 0.78 | 0.69 | 0.55 | 0.62 |

The resulting pair-wise similarity score for organisms A and B based on reaction R01 is *S*_*R*01_(*A, B*) = 0.87, and in general *S*_*R*_(*A, B*) is calculated for each of the shared reactions as shown in Tab. 5. Reactions, which are present in one organism but not the compared strain, offer the possibility for different scores to include the reaction, a scheme of the three different variants (one-to-one, one-to-many, and one-to-non) is illustrated in Fig. 8.

**Fig 8.**
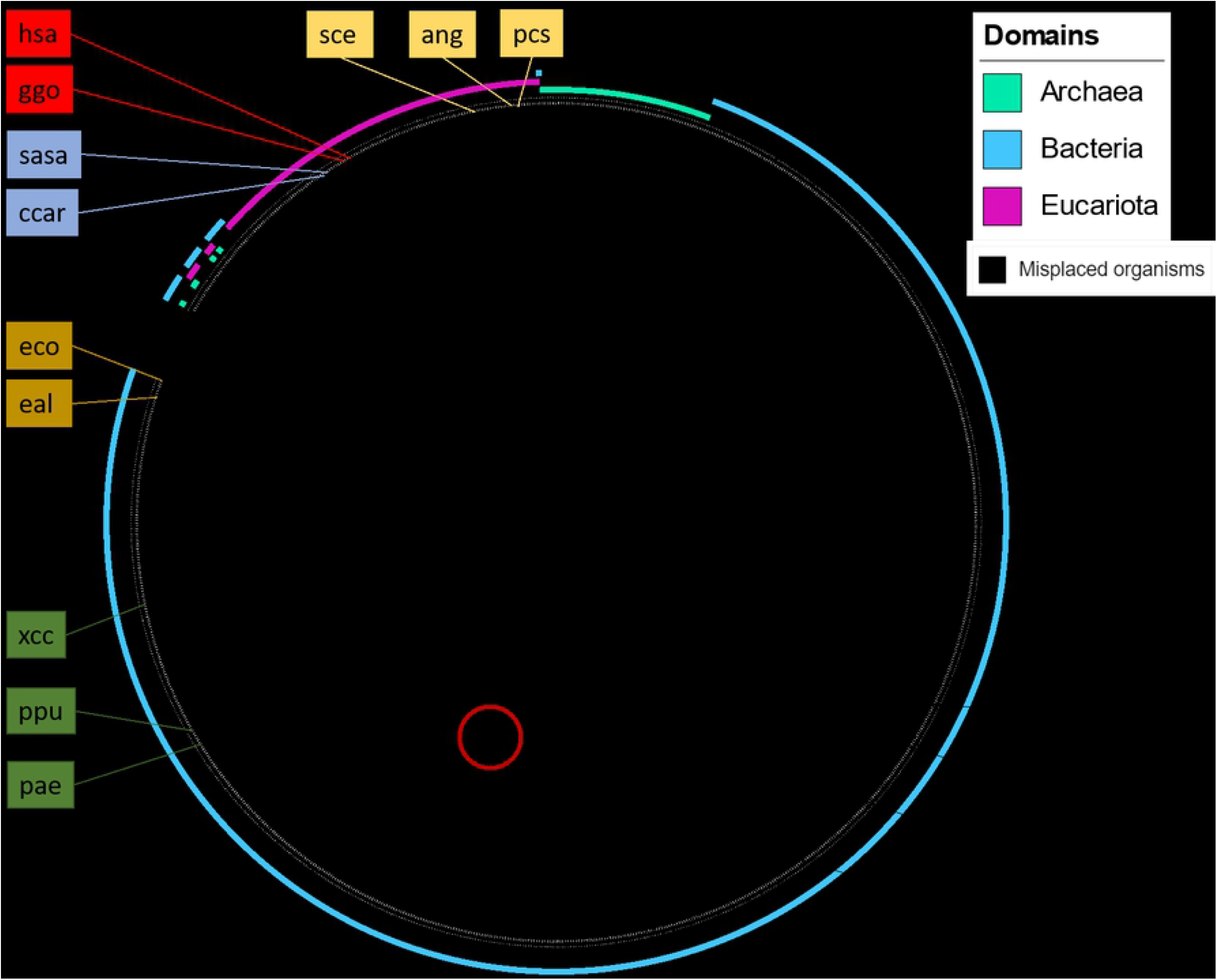
Four different reactions (R01-R04) for three different organisms (A-C) are compared. In reaction R01, a standard comparison with one enzyme in each organism is shown. Reaction R02 is not present in organism B, resulting in replacing the score with the penalty. Reactions R03 and R04 show the possibility of multiple proteins for one reaction (complexes and isozymes) and with the reaction missing in organism B, respectively. Below each case we show the resulting scores for the respective reaction comparisons.

In Fig. 8, reaction R01 illustrates the simplest case of comparisons, where each of the three organisms A, B, and C contain R01, which is encoded with a single enzyme. In contrast for reactions R02 and R04, organism B does not contain this metabolic capability, which excludes a sequence alignment. This is also shown in Tab. 5 with an entry of value X. We suggest that an excluded reaction can be accounted for in the functional similarity scoring in two ways: First, ignoring this reaction for the comparison, thus truly excluding it. Second, setting the comparison score to a scalar value serving as a possible penalty for this reaction comparison. Tab. 7 shows the consequence of different penalty values on the example using the data in Tab. 5. The reactions R03 and R04 in Fig. 8 show the case of multiple sequences for one metabolic function as previously described.

**Table 7.** Consequence of the penalty scores in combination with the example comparison of organism A and B in Tab. 5. Note that *J* (*A, B*) = 0.5 as a contrast.

| Penalty value | -0.25 | -0.1 | 0 | 0.1 | 0.25 | ignore penalty |
| --- | --- | --- | --- | --- | --- | --- |
| $\langle S(A, B) \rangle$ | 0.305 | 0.38 | 0.43 | 0.48 | 0.55 | 0.86 |

### Organisms selected for comparison

The 975 organisms used in the comparisons were manually selected amongst Eukaryota, Archaea, and Bacteria based on availability and distribution in the KEGG database. Organism families with many members, e.g. *E. coli* has 65 different strains stored in the KEGG database, were represented with multiple entries, whereas a strain occurring once might not be selected. This method of including organisms was chosen to evaluate the correct groupings of species, especially with regards to close strains compared to distant strains. A complete listing of the selected organisms’ KEGG IDs can be found in Supplementary Table 1.

The further subset of 21 organisms was selected in sets of threes, where two organisms are considered closer related to each other than to the third one based on standard sequence alignments, as shown in Tab. 4. This is illustrated by *H. sapiens, G. gorilla*, and *O. orca*: all three are mammals, two are closer related to each other than to the third one. Thus, the 21 organisms cover a large taxa range in the Tree of Life, while at the same time some organisms are very closely related. In Fig. 3, we have highlighted 16 of the 21 organisms within the tree of life to illustrate the coverage and distribution of the selected set of 21 organisms. The groupings chosen of different taxa are color coded consistently throughout the manuscript.

The access to the KEGG database using AutoKEGGRec to generate the genome-scale metabolic networks and extracting the protein sequences from KEGG was conducted during August 2018.

## Supporting information

**Supplementary table 1 Reactions of glycolysis**

**Supplementary table 2 Reactions of the pyruvate metabolism**

**Supplementary table 3 Reactions of the DNA metabolism**

**Supplementary table 4 Reactions of the TCA cycle**

**Supplementary table 5 Listing of reactions shared among all of the 21 organisms**

**Supplementary table 6 Listing of reactions used in at least 925 organisms**.

**Supplementary table 7 Listing of 975 organisms**

**Supplementary table 8 Jaccard-Index for misplaced organisms**

**Supplementary figure 1 Side by side comparison of Figure 4 and 5 A**

## Acknowledgments

EA and CS would like to thank ERA-IB-2 and The Norwegian Research Council grant #271585.

## Declarations

The funders had no role in study design, data collection and analysis, decision to publish, or preparation of the manuscript.

The authors have declared that no competing interests exist.

